# An integrated atlas of RNA to protein concordance across human tissues and cell types

**DOI:** 10.64898/2026.07.13.738336

**Authors:** Caitlin A. Finney, Artur Shvetcov

## Abstract

Differential expression studies, single-cell atlases, and target discovery pipelines measure RNA as a proxy for protein. Across human tissues, RNA explains only 40 to 60% of the variance in protein, and no framework resolves which genes reliably translate RNA into protein. We present an integrated atlas that matches single-cell transcriptomics from Tabula Sapiens and the Human Brain Cell Atlas to cell type-resolved immunohistochemistry from the Human Protein Atlas, comprising 488,190 observations across 11,154 genes, 24 tissues, and 53 cell types. For each gene we defined a suppression rate and separated genes into concordant, variable, and suppressed classes. The pooled correlation of ρ ≈ 0.4 reflects the mixing of these classes, and concordance depends on the interaction between gene and tissue. Suppression is predictable from gene sequence alone and traces to reduced translational efficiency and assembly-dependent degradation rather than to mRNA decay. The suppressed class is enriched for drug targets nominated on clinical evidence but not those validated by compound activity. We implement these classifications in an R package, concordR, and audit proteins nominated as brain-derived targets in neurodegeneration, where almost none survive at the protein level. Our atlas establishes RNA to protein concordance as a measurable property of the individual gene.

## Introduction

Differential expression studies, single-cell atlases, and mechanistic and target discovery pipelines all measure RNA and treat its expression as a proxy for protein. Here, RNA is used to assign function to genes, attribute activity to cell types, infer disease mechanisms, and nominate therapeutic targets. Across human tissues, however, RNA explains only 40 to 60% of the variance in steady state protein levels ^1–4^. The remaining variance reflects miRNA-mediated repression ^5–7^, codon optimality-coupled mRNA decay ^8,9^, stoichiometric degradation of orphan complex subunits ^10,11^, and protein half-lives that span five orders of magnitude ^1,12^. These regulatory processes are individually well characterized ^2,13^, yet no framework resolves which genes reliably translate their RNA into protein across human tissues and which do not.

The RNA to protein relationship is typically summarized by a single global correlation, which sits near ρ = 0.4 in organisms from yeast to humans ^2,14,15^. This single value is misleading, however, because it largely reflects differences between genes rather than RNA tracking protein within a gene. Each gene carries a characteristic RNA to protein ratio that is stable across tissues but varies by orders of magnitude between genes ^16^. When variation between genes is separated from variation of the same gene across tissues, scaled RNA accounts for the between-gene component but not the within-gene component ^17^. Matched RNA and protein have been measured at scale across human tissues ^18,19^, and protein to RNA ratios in those data have been predicted from sequence ^20^. These resources measure bulk tissue, and they neither separate genes by how reliably protein follows RNA nor resolve the cell types in which each gene is expressed. Single-cell transcriptomics has been paired with antibody-based protein profiling to classify the tissue specificity of human genes ^21^, but not to test RNA to protein concordance. A pooled correlation therefore cannot separate genes whose protein follows their RNA from genes whose protein does not, and this separation has not been made at the level of the individual gene across human cell types.

Here, we present an integrated RNA to protein atlas that matches single-cell transcriptomics from Tabula Sapiens ^22^ and the Human Brain Cell Atlas ^23^ to cell type-resolved immunohistochemistry (IHC) from the Human Protein Atlas ^24^. The resulting atlas comprises 488,190 matched observations across 11,154 genes, 24 tissues, and 53 cell types, with molecular and sequence features computed for every gene. For each gene, we defined a suppression rate, the fraction of high RNA observations in which protein went undetected, and used it to separate genes into concordant, variable, and suppressed classes. We show that the pooled correlation of ρ ≈ 0.4 reflects the mixing of these classes and that concordance is a property of the interaction between gene and tissue. We further show that suppression can be reasonably predicted from gene sequence alone and traces to reduced translational efficiency and assembly-dependent degradation rather than to mRNA decay. The suppressed class is enriched for drug targets nominated on clinical and literature evidence but not for targets validated by direct compound activity, highlighting that RNA-based target nomination is unreliable for these genes. We implement these classifications in an R package, concordR, and use it to audit proteins nominated as brain-derived mechanisms and drug targets in neurodegeneration. Across published cerebrospinal fluid and plasma studies, almost none of these nominations can be supported at the protein level. Our atlas establishes RNA to protein concordance as a measurable property of the individual gene and provides an open resource for testing it across tissues and cell types (Fig. 1a).

**Figure 1.**
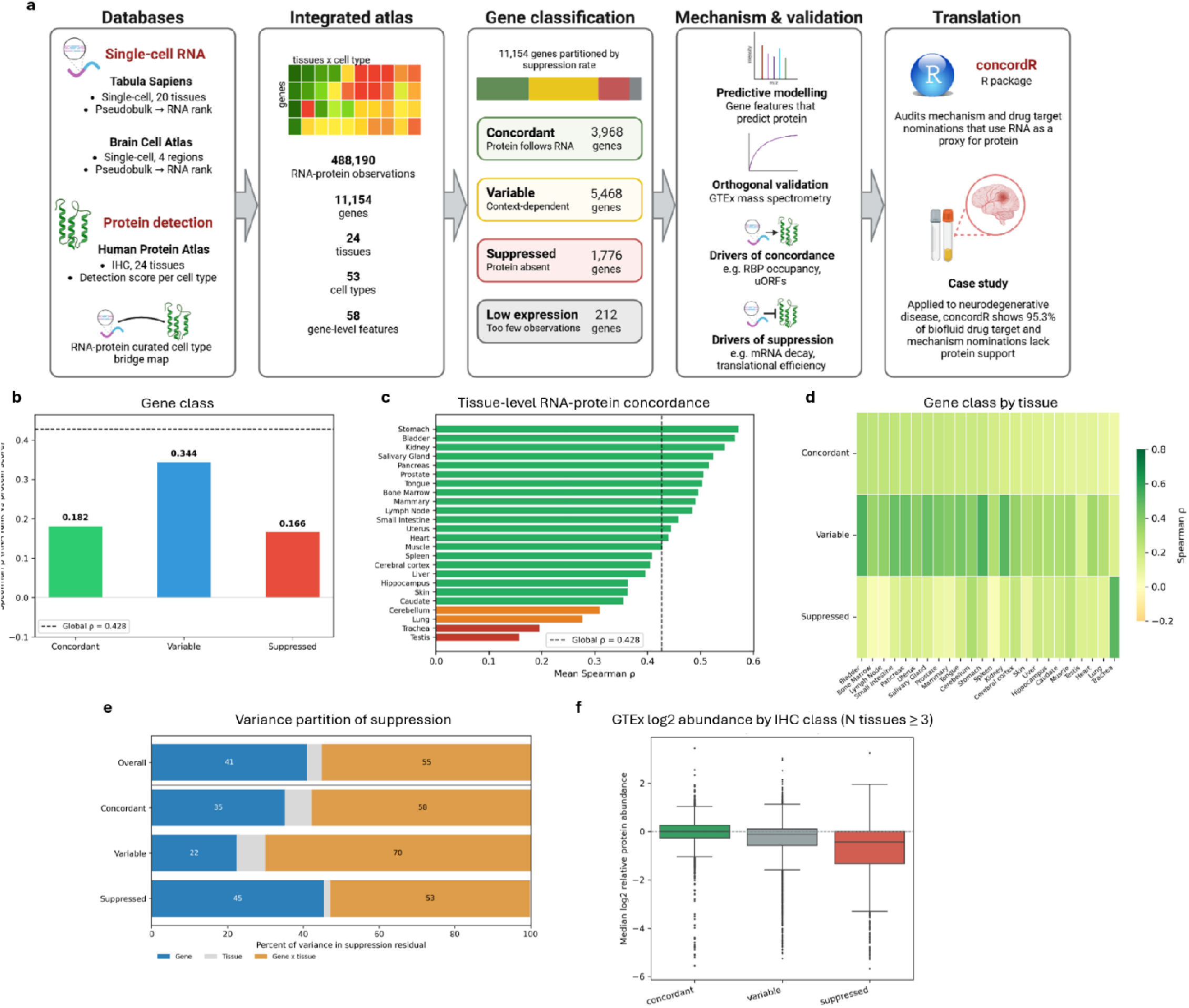
An integrated atlas resolves RNA to protein concordance and shows the pooled correlation is inflated by mixing gene classes. (a) Schematic of atlas construction and the current study. (b) Spearman correlation between RNA rank and protein detection, computed globally and within each gene class. The global value was ρ = 0.428. Every within-class correlation fell below it (concordant ρ = 0.182, variable ρ = 0.344, suppressed ρ = 0.166). (c) Spearman correlation within each of the 24 tissues. Values ranged from ρ = 0.572 in stomach to ρ = 0.145 in testis. (d) Spearman correlation within each gene class across the 24 tissues. Variable genes showed the highest correlation in 22 of 24 tissues (mean ρ = 0.363, SD = 0.05). Concordant and suppressed genes were lower (mean ρ = 0.178, SD = 0.106 and mean ρ = 0.162, SD = 0.101). (e) Variance in per-gene discordance partitioned into gene identity, tissue, and their interaction, shown across all genes and within each class. Across all genes, gene identity accounted for 41% of the variance, tissue for 4%, and their interaction for 55%. Within the suppressed class, gene by tissue interaction carried the most weight at 53% and tissue the least at 1.8%. Within the variable class, the gene by tissue interaction dominated at 70%. (f) Median protein abundance from the GTEx TMT-MS3 proteome across the three IHC-defined gene classes. Abundance decreased from concordant through variable to suppressed genes (Cliff’s δ = 0.41 concordant versus suppressed, Kruskal-Wallis p < 0.0001, pairwise Mann-Whitney adjusted p < 0.0001).

## Results

### An integrated atlas resolves transcript abundance and protein detection at cell type resolution

To quantify the relationship between transcript abundance and protein levels, we constructed an integrated atlas in which both are measured in matched tissues and cell types. Single-cell RNA sequencing data from Tabula Sapiens ^22^ (Spearman’s ρ = 0.436, n = 410,430) and the Brain Cell Atlas ^23,25^ (Spearman’s ρ = 0.383, n = 77,760) were aggregated into pseudobulk expression profiles and harmonized against immunohistochemistry (IHC) protein scores from the Human Protein Atlas ^24^ using a manually curated tissue harmonization map. The brain was split into five anatomical regions across the caudate, cerebellum, cerebral cortex, hippocampus, and hypothalamus. However, given the low number of genes detected (<100) hypothalamus was removed prior to further analyses. After filtering based on antibody reliability, detection level, and a minimum of 50 contributing cells, the atlas comprises 488,190 observations spanning 11,154 genes, 24 tissues, and 53 cell types. For each observation we computed 58 gene level features spanning protein biophysics, degradation motifs, transcript structure, translational efficiency, subcellular localization, complex membership, microRNA targeting, upstream open reading frames, and RNA binding protein occupancy. Across the complete resource, the global RNA to protein correlation was ρ = 0.428. This is consistent with previous reports of moderate RNA to protein correlation ^1–3^.

### The pooled RNA to protein correlation is inflated by mixing gene classes

Despite this low-moderate correlation, many still rely on using RNA as a proxy for protein. A single value, however, masks variation among genes ^1,16,18^, so we next sought to determine whether this correlation reflects a property that genes share, or an artefact of pooling subsets of genes with different translational fate. We first computed a suppression rate for each of the 11,154 genes utilizing the fact that every gene appears in the atlas several times, across different tissues and cell types. Suppression rate was defined as the fraction of a gene’s transcript observations in which protein went undetected, ranging from reliably detected to absent. Within this framework, we labelled genes as ‘concordant’ whose protein reliably follows their RNA (suppression rate <0.2; n=3,698), ‘suppressed’, whose protein is largely absent despite abundant RNA (suppression rate >0.8, n=1,776), and ‘variable’, whose protein presence largely depends on context (>0.2 and <0.8; n=5,468). A further 212 genes had insufficient transcript observations to score and were excluded as low expression. In every class the within-class correlation fell below the pooled value of ρ = 0.428 (Fig. 1b). For concordant and suppressed genes this was expected as both classes are defined by protein detection irrespective of RNA (ρ = 0.182 and 0.166; Fig. 1b). Importantly, although higher than in other groups, the variable group was still below the global value (ρ = 0.344; Fig. 1b) despite being the only class in which RNA carries genuine information about protein.

We then sought to determine if this finding extends across tissues and cell types. Tissue level correlations were highly variable, ranging from ρ = 0.572 (stomach) to ρ = 0.145 (testis) (Fig. 1c). Within each gene class, however, the rank order was largely comparable across tissues. Variable genes consistently showed the highest correlation (mean ρ = 0.363, SD=0.05; 22/24 tissues; Fig. 1d). Concordant and suppressed genes were both lower (mean ρ = 0.178, SD=0.106 and ρ = 0.162, SD=0.101, respectively; Fig. 1d). Highly variable correlations were also found at the cellular level (Supplementary Fig. 1), ranging from ρ = 0.572 in stomach epithelial cells down to ρ = 0.077 in testis peritubular myoid cells. These findings suggest that tissue-level concordance is therefore organized by the gene class rather than by an intrinsic property of the tissue or cell type themselves.

To confirm this, we partitioned the variance in per-gene discordance into three parts, that attributable to gene identity, to tissue, or that depends on the specific combination of the two. Overall, gene identity accounted for 41% of the variation, tissue for only 4%, with the remaining 55% explained by the gene-by-tissue combination (Fig. 1e). The same partitioning within each gene class confirmed that all three reflect different regulatory scenarios. Here, gene by tissue interaction carried the most weight (53%) and tissue the least (1.8%) in suppressed genes (Fig. 1e). Variance in variable genes, however, was dominated by the gene-by-tissue combination (70%). Therefore, the RNA-protein concordance within a particular tissue does not reflect intrinsic properties of the tissues. Instead, it reflects the interaction of genes within tissues and the resulting context-dependent behavior of variable genes.

To establish that the concordance classes reflect genuine protein abundance rather than antibody performance, we compared them against the GTEx TMT-MS3 proteome (12,627 genes, 15 tissues) ^18^, a platform that shares no methodology with IHC beyond the protein as a target. Thus, agreement between the two cannot arise from shared technical artefacts. We found that the median MS protein abundance decreased monotonically across the IHC-defined classes (concordant > variable > suppressed), with a large separation between the extremes (Cliff’s δ = 0.41, concordant vs suppressed; Kruskal-Wallis *p* < 0.0001; pairwise Mann-Whitney adjusted p < 0.0001) (Fig. 1f). This ordering was robust across detection coverage thresholds and reproduced in continuous form (protein confidence vs MS abundance, Spearman’s ρ = 0.268). Finally, because immunohistochemistry can miss proteins that actively leave the cell, we tested secretion directly as a confounder on TMT-MS. Secreted suppressed-class genes were not recovered but showed the lowest abundance of any group (median -0.69 vs -0.27 for non-secreted suppressed genes and 0.00 for concordant genes; δ = -0.13, p-value < 0.001), excluding antibody-export artefact as the basis of the suppressed class. This consistently clear separation on the MS orthogonal abundance confirms that these RNA-protein discordance findings reflect genuine protein output rather than an artefact of IHC.

### Gene level sequence and molecular features predict protein detection independently of RNA abundance

Having shown that RNA poorly correlates with protein for most genes, we asked whether this discordance is determined or stochastic. We therefore tested whether protein detection can be predicted from gene features alone. Here, determined discordance would be reflected by an ability of gene features to predict protein whereas if the discordance is stochastic, they would not. To test this, we trained gradient boosted classifiers to predict protein detection, under five-fold cross validation grouped by gene so that no gene appeared in both the training and test sets. We compared models built from the full set of 58 gene level features other than RNA rank, RNA rank only, or 59 features (gene level features + RNA rank). We found that the model with 58 gene features reached AUC=0.755 (Fig. 2a). A model using RNA expression rank only had a similar ability to predict protein detection (AUC=0.749; Fig. 2a). A model combining RNA rank and the gene features led to an improvement in the AUC to 0.779 (95% CI 0.7742-0.7844; Fig. 2a), with well calibrated predicted probabilities across the full range (Brier score = 0.1858, ECE = 0.0081). Further, model performance was not influenced by the specific dataset, with gene level protein confidence highly correlated across Tabula Sapiens and Brain Cell Atlas (ρ = 0.7772; Fig. 2b). Extending this, training on Tabula Sapiens (410,430 observations, 20 tissues) and testing on the Brain Cell Atlas (77,760 observations, 4 regions) produced similar AUCs (0.722-0.816) across model types (Fig. 2c). To determine which variables drove performance in our full model (58 gene level features and RNA expression rank) we used permutation importance. Here, RNA rank was identified as the largest contributor (ΔAUC = 0.126; Fig. 2d). Permutation importance of the gene features only model indicated that RBP occupancy was the dominant feature outside of RNA rank (ΔAUC = 0.035; Fig. 2e).

**Figure 2.**
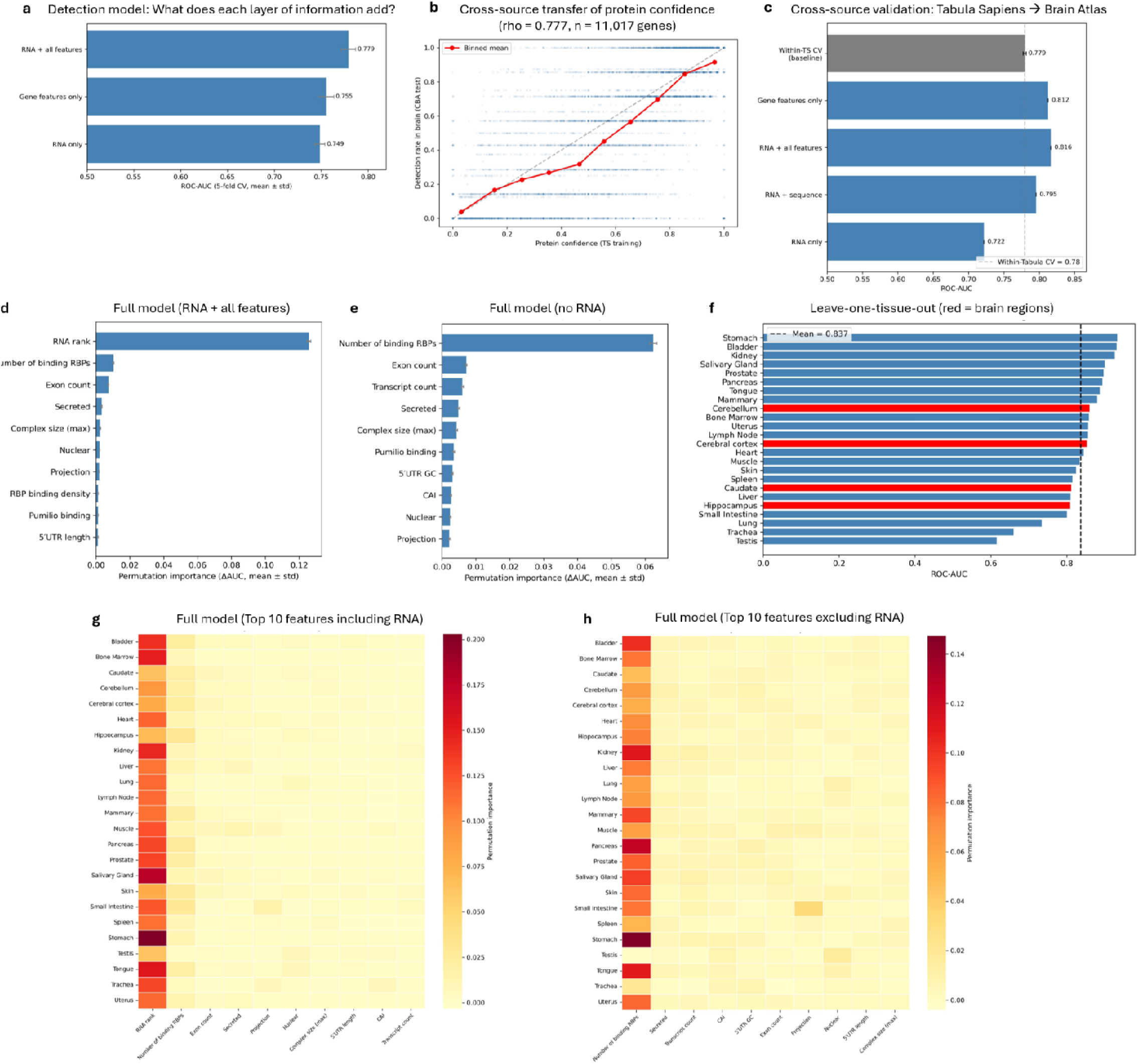
Gene sequence and molecular features predict protein detection independently of RNA. (**a**) ROC-AUC for three gradient-boosted models predicting protein detection under gene-grouped five-fold cross-validation, with no gene shared between training and test sets. The 58 gene features reached AUC = 0.755. RNA rank alone reached AUC = 0.749. The combined model reached AUC = 0.779 (95% CI 0.7742 to 0.7844), with well-calibrated predictions (Brier score = 0.1858, ECE = 0.0081). (**b**) Gene-level protein confidence correlated across Tabula Sapiens and the Brain Cell Atlas (Spearman ρ = 0.7772). (**c**) ROC-AUC for models trained on Tabula Sapiens (410,430 observations, 20 tissues) and tested on the Brain Cell Atlas (77,760 observations, 4 regions). AUCs ranged from 0.722 to 0.816 across model types. (**d**) Permutation importance for the full model of 58 gene features and RNA rank. RNA rank was the largest contributor (ΔAUC = 0.126). (**e**) Permutation importance for the gene-features model. RNA-binding protein (RBP) occupancy was the largest contributor (ΔAUC = 0.035). (**f**) ROC-AUC for the full model under leave-one-tissue-out evaluation, training on 23 tissues and testing on the held-out tissue. The mean AUC was 0.834, and performance across the four brain regions was 0.8162. Performance was lowest in testis (AUC = 0.611) and trachea (AUC = 0.642). (**g**) Permutation importance for the full model across tissues. RNA rank was again the largest contributor (mean ΔAUC = 0.12). (**h**) Permutation importance for the gene-features model across tissues. RBP occupancy led in 23 of 24 tissues (mean ΔAUC = 0.060). In testis, nuclear localization dominated (ΔAUC = 0.020).

We then evaluated whether the full model was generalizable across tissues. To do this, we trained gradient boosted classifiers using a leave-one-tissue-out evaluation, training on 23 tissues and testing on the held-out tissue. The model predicted protein detection in the held-out tissue with a mean AUC of 0.834, and performance across the four brain regions was similar at 0.8162 (Fig. 2f). It was lowest in testis (AUC 0.611) and trachea (0.642), the former reflecting the specialized post-meiotic programme of testis ^26,27^ and the latter likely underpowered (n = 5,130). Permutation importance analysis of the full model across tissues indicated that the importance of RNA expression rank was again the largest contributor (mean ΔAUC=0.12; Fig. 2g). Similarly, in the gene features only model, RBP occupancy was the leading feature in 23 of 24 tissues (mean ΔAUC=0.060; Fig. 2h). In testis, nuclear localization (ΔAUC = 0.020) dominated, consistent with the specialized post-transcriptional program of spermatogenesis, in which canonical miRNA-mediated regulation is attenuated ^28,29^.

### Concordant and suppressed genes are distinguished by multi-feature structures

Although our models identified the gene level features that predict protein abundance, the direction of the effects and the features (molecular mechanisms) underlying the ability to separate reliably detected from suppressed genes remains unknown. To resolve this, we performed a systematic comparison across all 58 gene level features using Cohen’s d. For concordant genes, total bound RBPs per transcript was the dominant discriminator (d = 1.5; *p* < 0.0001), followed by Pumilio binding (d = 1.3; *p* < 0.0001), HuR/ELAVL1 binding (d = 0.94; *p* < 0.0001), HNRNP binding (d = 0.82; *p* < 0.0001), and uORF presence (d = 0.81; *p* < 0.0001) (Fig. 3a). uORFs in concordant genes were also longer (mean 21.4 vs. 9 amino acids; d = 0.65; *p* < 0.0001) and more conserved (mean PhastCons 0.32 vs. 0.13; d = 0.63; *p* < 0.0001), suggesting uORFs are playing a role in fine tuning protein output rather than acting as full repressors. Other features associated with concordance included nuclear localization (d = 0.63; *p* < 0.0001), protein-complex membership (large-complex membership d = 0.52; p-value < 0.0001), and transcript count (d = 0.45; p-value < 0.0001; Fig 3a). In contrast, suppression was characterized by a weaker signature, with the secreted protein as the strongest feature (d = -0.75; *p* < 0.0001; Fig. 3a). As shown above, this finding is not due to a bias against secreted proteins in IHC. The remaining features associated with suppression included N-terminal hydrophobicity (d = -0.43; p-value < 0.0001), N-terminal hydrophobic run length (d = -0.41; *p* < 0.0001), and cysteine fraction (d = -0.39; *p* < 0.0001; Fig. 3a).

**Figure 3.**
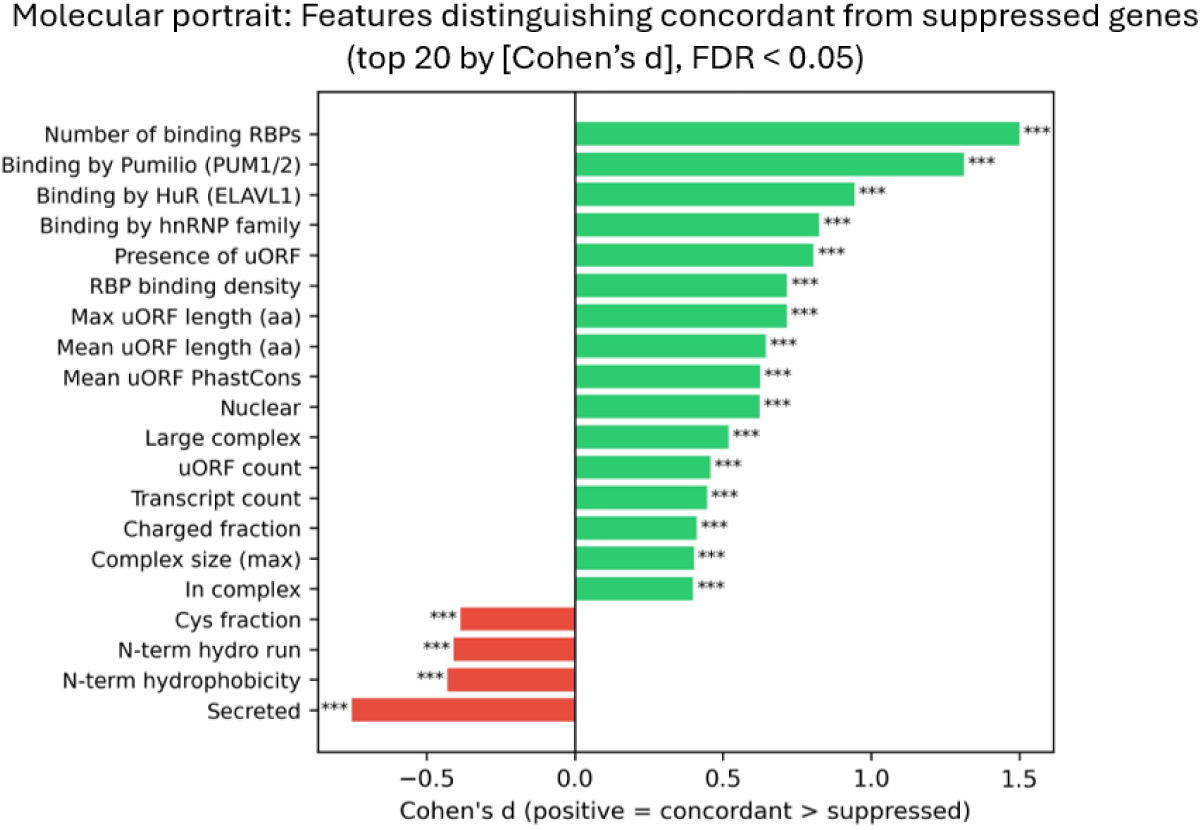
Concordant and suppressed genes are distinguished by multi-feature structures. Effect size (Cohen’s d) for each of the 58 gene features comparing concordant and suppressed genes. Positive values mark features higher in concordant genes. Negative values mark features higher in suppressed genes. The strongest concordant features were total bound RBPs per transcript (d = 1.5), Pumilio binding (d = 1.3), HuR/ELAVL1 binding (d = 0.94), HNRNP binding (d = 0.82), and uORF presence (d = 0.81). The strongest suppressed feature was the secreted protein annotation (d = -0.75). All reported features passed p < 0.0001.

We next sought to clarify whether the multi-feature structure of protein detection is determined additively, through the sum of many modest contributions, or through more complex interactions across features. To test this, we compared AUCs from a logistic regression, which is only additive, and a gradient-boosted classifier, which accounts for interactions between features, both of which were fitted to the concordant and suppressed population on the 15 highest effect size features. The two performed almost identically (logistic AUC = 0.845 ± 0.014; nonlinear AUC = 0.849 ± 0.010), indicating that an additive model is sufficient to separate the gene classes and little additional signal is carried by feature interactions.

These data highlight that concordance is characterized by numerous features related to RBP and uORF mechanisms and stabilization through protein complex membership, whereas suppression is characterized by signal peptide and secretory pathways. Further, this multi-feature structure of protein detection is additive rather than the result of complex relationships between these features.

### Assembly-dependent degradation links binding partner availability to subunit detection

Among the dominant features distinguishing concordance from suppression was stabilization through protein complex membership. To determine the mechanisms underlying this relationship, we tested whether this was the result of rapid degradation of unassembled protein complex subunits. To do this, we computed the Spearman correlation between mean partner RNA rank and focal subunit detection for 6,716 gene-complex pairs, 1,502 CORUM v5.0 complexes across 24 tissues. Binding partner expression and subunit detection were positively correlated in 67.3% of gene-complex pairs (p < 0.0001) and 73.0% of complexes (p < 0.0001), though the effect was modest at ρ = 0.119 (Fig. 4a,b). We then checked whether this was reflective of assembly-dependent degradation rather than general co-expression among related genes by shuffling partner identities (500 permutations). This collapsed the correlation to a null mean of 0.019 ± 0.003, placing the observed value 33 standard deviations above the null (permutation p = 0.002; Fig. 4a). The positive correlation between binding partner expression and subunit detection also varied across the three gene classes from variable genes (ρ = 0.138) followed by concordant (ρ = 0.105) and suppressed genes (ρ = 0.078; Fig. 4c). Further, we found the largest effect sizes from obligate heteromers whose subunits are known to undergo ER-associated degradation, including GABA_A_ receptors (ρ = 0.70), NMDA receptors (ρ = 0.58-0.64), and SNARE complexes (ρ = 0.55-0.58; Fig. 4d).

**Figure 4.**
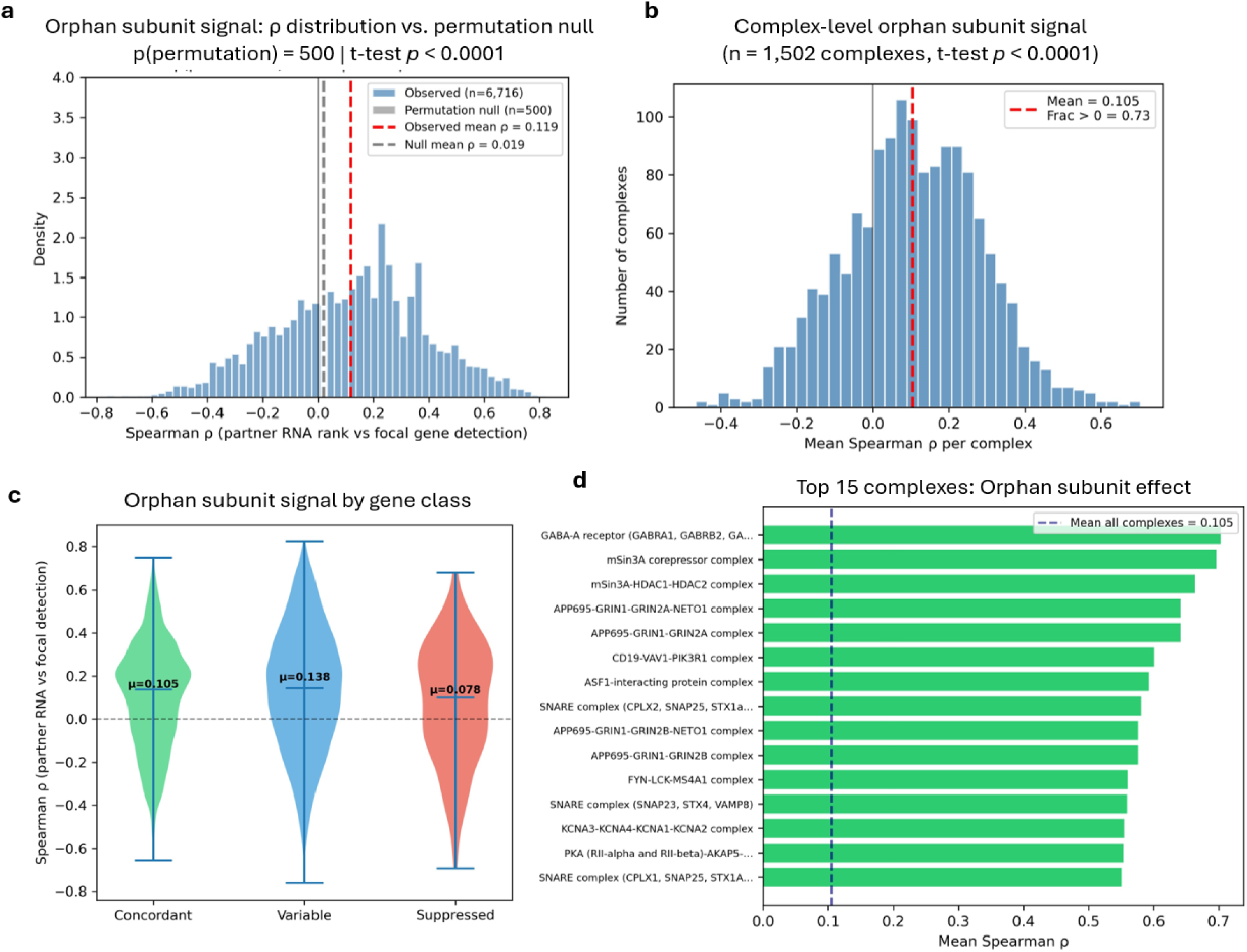
Assembly-dependent degradation links binding partner availability to subunit detection. (**a**) Distribution of per-pair Spearman correlations between mean partner RNA rank and focal subunit detection across 6,716 gene-complex pairs. The observed mean was ρ = 0.119. Partner expression and subunit detection were positively correlated in 67.3% of pairs (p < 0.0001). A permutation test shuffling partner identities 500 times collapsed the correlation to a null mean of 0.019 ± 0.003. This placed the observed value 6 standard deviations above the null (permutation p < 0.002). (**b**) Per-complex correlations across 1,502 CORUM v5.0 complexes. The correlation was positive in 73.0% of complexes (p < 0.0001). (**c**) Correlation between partner expression and subunit detection within each gene class. The effect was largest in variable genes (ρ = 0.138), followed by concordant genes (ρ = 0.105) and suppressed genes (ρ = 0.078). (**d**) Correlation for obligate heteromers whose subunits undergo ER-associated degradation. The largest effects were in GABA_A_ receptors (ρ = 0.70), NMDA receptors (ρ = 0.58 to 0.64), and SNARE complexes (ρ = 0.55 to 0.58).

Together these results identify assembly-dependent degradation as one mechanism separating concordant from suppressed genes. The effect depended on the specific partners of each subunit and not on shared regulation among related genes. Although it was small for any single subunit, it was detectable in the majority of complexes tested.

### Reduced translation is the primary route to RNA-protein suppression

To further determine and characterize when suppression occurs mechanistically, we compared suppressed and concordant genes using three measurements: mRNA half-life (RNADecayCafe ^30^), translational efficiency (TE) from ribosome profiling (RPFdb v3.0 ^31^), and protein half-life ^12^. Suppressed genes showed no shortening of mRNA half-life relative to concordant genes. In a cell type matched comparison there was no difference (d = -0.01, *p* = 0.82; Fig. 5a) and across all cell lines the small difference pointed in the opposite direction (median 3.22 versus 3.02 hours; d = -0.11, *p* = 0.005; Fig. 5b), excluding mRNA decay as the mechanism. In contrast, TE was reduced by approximately 50% in suppressed genes (median TE = 0.437 versus 0.916 in concordant; d = 0.71, *p* < 0.0001; Fig. 5c). Here, the TE distribution was uniformly shifted rather than bimodal, indicating graded translational attenuation across all suppressed genes rather than a discrete subpopulation. Protein half-life was modestly shorter in suppressed genes (median 55.7 versus 75.1 hours; d = 0.16; *p* = 0.0007; Fig. 5d). Of note, protein half-life was only measured for 15.8% of suppressed genes and biased toward hematopoietic and hepatocyte cell types. However, this is consistent with post-translational degradation acting as a secondary contributor that compounds the translational deficit rather than as an independent mechanism. We then cross-referenced the suppressed genes against orthogonal mass spectrometry evidence for 5′UTR micropeptide translation using the ReSpin dataset in sORFs.org ^32^. We found that only 14/1,776 suppressed genes (0.8%) carried high confidence evidence (>95%) of a translated micropeptide product.

**Figure 5.**
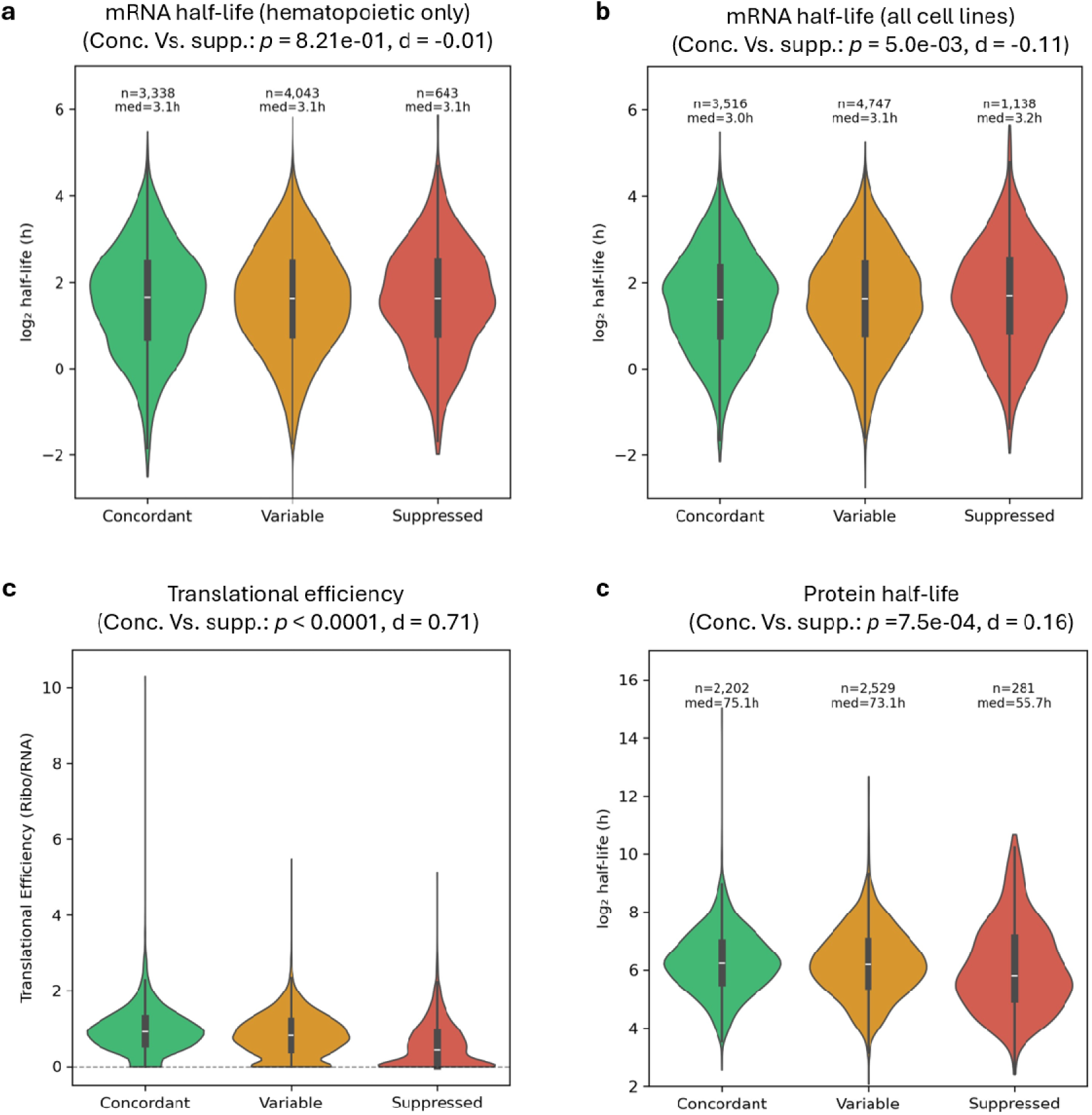
Reduced translation is the primary route to RNA-protein suppression. (**a**) mRNA half-life in a cell type matched comparison of suppressed and concordant genes. The two classes did not differ (Cohen’s d = -0.01, p = 0.82). (**b**) mRNA half-life across all cell lines. Suppressed genes showed a slightly longer median half-life than concordant genes (3.22 versus 3.02 hours, d = -0.11, p = 0.005). (**c**) Translational efficiency from ribosome profiling. Suppressed genes showed roughly half the efficiency of concordant genes (median 0.437 versus 0.916, d = 0.71, p < 0.0001). (**d**) Protein half-life. Suppressed genes showed a modestly shorter median half-life than concordant genes (55.7 versus 75.1 hours, d = 0.16, p = 0.0007).

Together these results identify reduced translational efficiency as the primary route to suppression, with mRNA decay excluded and shortened protein half-life acting only as a secondary contributor. Micropeptide translation carried detectable evidence in fewer than one percent of suppressed genes and therefore cannot account for the translational deficit, which we measure directly from ribosome profiling.

### Suppressed genes are enriched for drug targets in clinical but not assay-based resources

Having established the main mechanisms underlying suppression, we next asked whether this carries translational consequences. Specifically, RNA-based drug target nomination assumes that transcript abundance tracks protein and we have shown that this assumption is not supported for many genes. We therefore tested whether the suppressed class is enriched for therapeutic targets. Among single protein targets in ChEMBL ^33^, defined by measured compound activity, suppressed genes showed no enrichment (OR = 0.89, ns; Fig. 6). Targets curated in resources incorporating clinical and literature evidence, however, were enriched among suppressed genes. This included the Drug Gene Interaction Database (DGIdb) ^34^ (OR = 1.36, p < 0.001), including for approved therapeutics (OR = 1.28, p < 0.001), and in Open Targets ^35^ (OR = 1.30, p = 0.003; Fig. 6). The suppressed class contains secretory proteins including AFP. These proteins are exported rather than retained in the cell, therefore their low atlas score could reflect a limitation of *in situ* staining rather than genuinely reduced protein. We tested this possibility using mass spectrometry, which does not depend on staining intensity, and found that the expression of these genes remained low.

**Figure 6.**
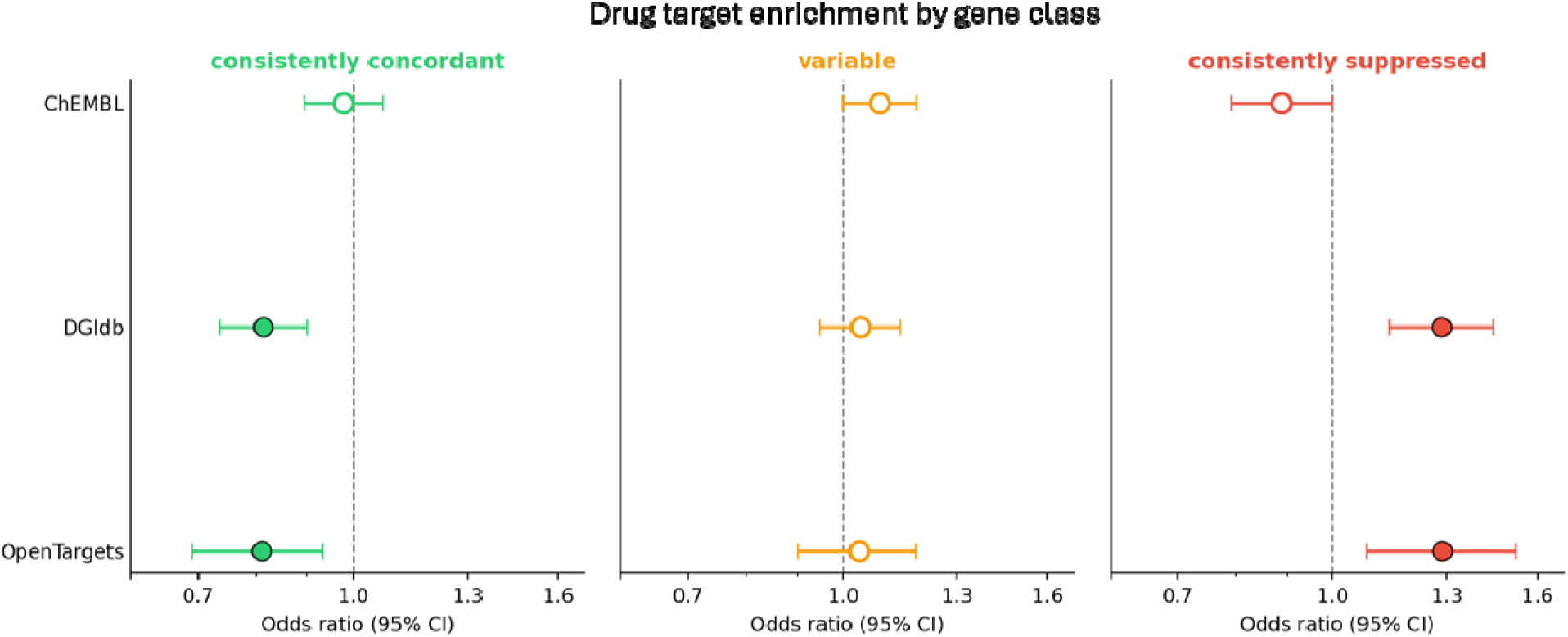
Suppressed genes are enriched for drug targets in clinical but not assay-based resources. Odds ratios for enrichment of the suppressed gene class across drug target resources. Each point shows the odds ratio and its 95% confidence interval, with the vertical line at an odds ratio of 1. Suppressed genes showed no enrichment among ChEMBL single protein targets defined by measured compound activity (OR = 0.89, ns). Suppressed genes were enriched among targets in the Drug Gene Interaction Database (OR = 1.36, p < 0.001), including approved therapeutics (OR = 1.28, p < 0.001), and among targets in Open Targets (OR = 1.30, p = 0.003).

The suppressed gene class is enriched for therapeutic targets in resources based on clinical and literature evidence but not in those based on measured compound activity. Although several of these targets are clinically established, their protein abundance is controlled after transcription rather than by it. Together, these findings highlight that we cannot reliably nominate drug targets within this class from RNA abundance alone.

### Most biofluid drug targets nominated in neurodegeneration cannot be attributed to brain

We have shown that abundant RNA does not guarantee detectable protein, and that this suppression is systematic. The suppressed class is also enriched for drug targets nominated from clinical evidence, so RNA-based nomination is unreliable for these genes. Despite this, researchers rely on RNA evidence in diseases where tissue is inaccessible, such as neurodegeneration where brain is sampled only at autopsy. In these studies, a protein found at differential abundance in cerebrospinal fluid (CSF), for example, is cross-referenced against an RNA atlas, reported as enriched in a brain cell type, and nominated as a brain-derived target. This treats one measure of RNA enrichment as evidence for the presence of the protein, its location, and its tissue of origin at once. To support the use of our atlas in translational research, we developed concordR (Methods) and used it to audit 3,304 CSF proteins nominated as brain cell type-enriched across published neurodegenerative disease studies ^36–44^ (Supplementary Table 1). Each protein was assessed against four criteria, (1) detection of protein in brain, (2) a physical route from brain to biofluid, (3) specificity of the protein to brain, and (4) concordance between RNA and protein. A protein counted as detected when supported by HPA immunohistochemistry, GTEx mass spectrometry, or PaxDb mass spectrometry, so a tissue mismatch required its absence across all three platforms. A route counted as plausible when the protein enters the biofluid through the secretory pathway or proteolytic cleavage. Neuronal projections release their contents into the extracellular space through synaptic turnover and axonal degeneration, so a route also counted as plausible for an intracellular protein carrying a neuronal projection annotation (e.g. NEFL).

### Assessed independently, each criterion removed a different share of the CSF proteins

Most proteins were detectable in brain (91.4%, 3,019/3,304; Fig. 7a), the exceptions being proteins whose RNA is present but whose protein lies elsewhere, such as CRYBB2, confined to the eye, and NUDT5, ubiquitous in RNA yet absent from brain at the protein level. The healthy donor atlases may not represent the diseased brain, so we compared the undetected proteins against an independent Alzheimer’s disease cohort (Accelerating Medicines Partnerships-Alzheimer’s Disease (AMP-AD); n = 1,275 donors ^45^) that underlies most of the studies audited, and only three further proteins were recovered (3,022/3,304, 91.5%), so disease state does not account for the absences. The remaining criteria flagged far more CSF proteins. A physical route to the biofluid was plausible for 56.6% (1,869/3,304), concordance between RNA and protein held for 21.0% (694/3,304), and specificity to brain for only 9.9% (328/3,304; Fig. 7a).

**Figure 7.**
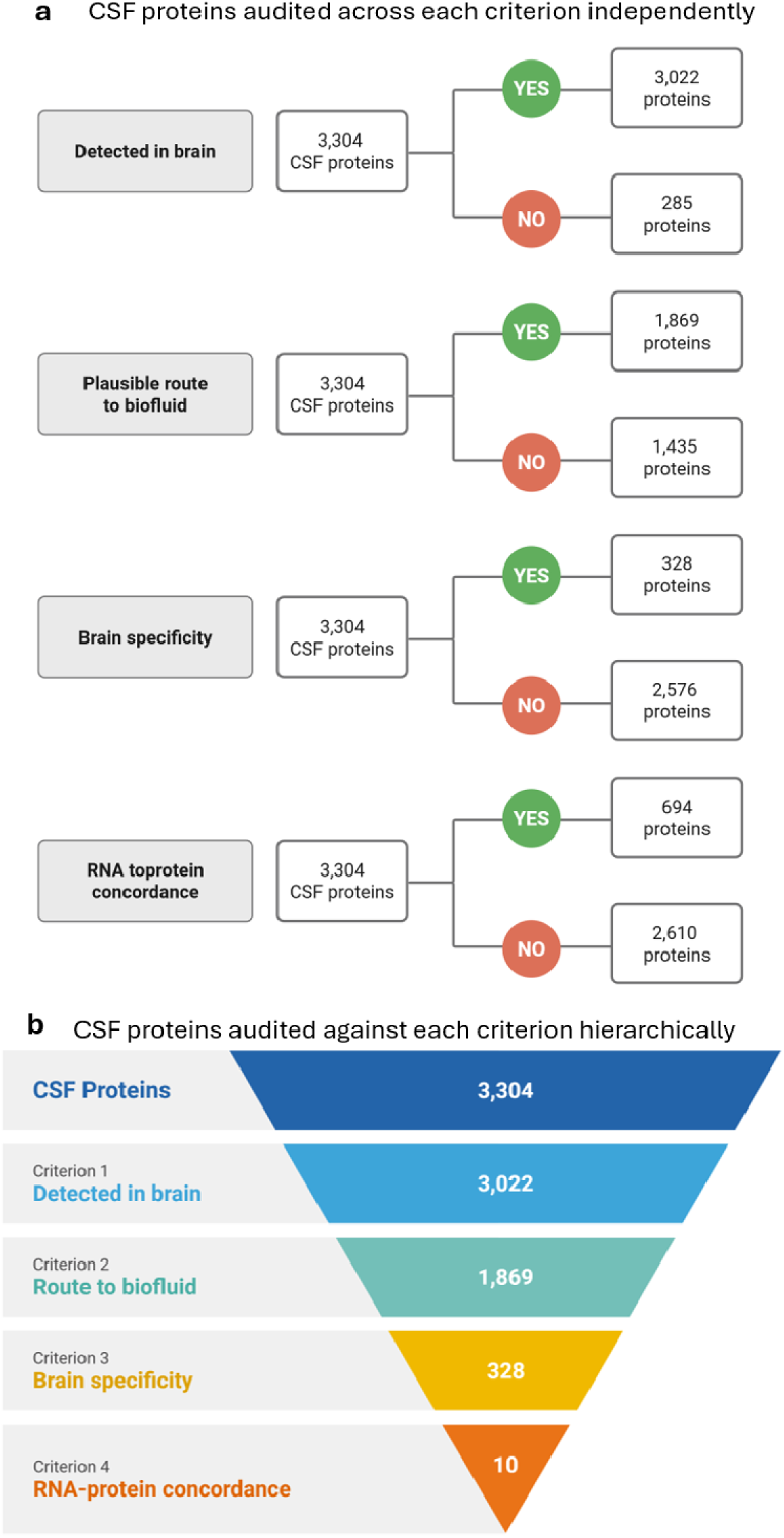
Most biofluid mechanism and drug targets nominated in neurodegeneration cannot be attributed to brain. (**a**) Share of 3,304 cerebrospinal fluid (CSF) proteins (see Supplementary Table 1 for full list) passing each of the four audit criteria assessed independently. Most proteins were detectable in brain (91.5%, 3,022/3,304). A physical route to the biofluid was plausible for 56.6% (1,869/3,304). RNA to protein concordance held for 21.0% (694/3,304). Specificity to brain held for 9.9% (328/3,304). (**b**) The same four criteria applied in sequence, shown as the number of proteins surviving each step. Of 3,304 CSF proteins, 3,022 were detected in brain, 1,869 had a plausible physical route to the biofluid, 328 were specific to brain, and 10 were concordant between RNA and protein. Ten proteins survived the full audit, a little over 0.3% of those nominated.

Applied in sequence, the criteria compound to a much smaller set. Of the 3,304 CSF proteins, 3,022 were detected in brain (Fig. 7b). Lumbar CSF draws roughly 80% of its proteins from blood, so brain detection alone cannot establish brain origin ^46–48^. Requiring a plausible physical route from brain to biofluid left 1,869, of which 328 were specific to brain and 10 were concordant between RNA and protein. Of every protein nominated in this literature as a drug target or mechanistic candidate in neurodegeneration, 10 survive as defensible, or a little over 0.3%.

Together, these findings show that almost none of the targets nominated from brain RNA in this literature can be defended at the protein level. RNA enrichment alone cannot establish that a biofluid protein comes from the brain, and without that evidence these nominations do not support target discovery.

## Discussion

The RNA to protein correlation sits near ρ = 0.4 across organisms and platforms ^2,3,14^. In the original yeast measurement it was 0.1 to 0.4 for most genes and reached 0.94 only when the eleven most abundant proteins were added ^14^. We show that pooling gene classes produces the same inflation. The correlation is a composition-weighted average of gene classes that convert RNA to protein with different reliability. Every within-class correlation in our atlas falls below the pooled ρ = 0.428, including the concordant class in which RNA carries reliable information about protein. A global correlation therefore cannot report whether RNA tracks protein for a given gene, and the reliability of RNA as a proxy must be assessed gene by gene.

Prior work has shown that RNA recovers differences between genes but not the variation of one gene across tissues ^2,3,17^, and that each gene carries its own RNA to protein ratio that holds across tissues ^16^. These results, however, describe the amount of protein made per unit of RNA, and all come from bulk tissue. They do not address whether a gene’s protein is present at all when its RNA is high, nor whether there are differences between the cell types of a tissue. Single-cell transcriptomics and antibody-based protein profiling have been paired to classify the tissue specificity of genes ^21^, but not to test concordance. We introduce a suppression rate, the fraction of a gene’s transcript observations in which protein goes undetected, computed for each gene across tissues and cell types. The variance in suppression rate partitions into gene identity (41%), tissue (4%), and their interaction (55%), establishing suppression as a property of the gene that varies by tissue and cell types.

In our atlas the suppressed class is defined from IHC. Undetected protein could therefore reflect antibody failure rather than genuine absence, or the class could be stochastic rather than a real feature of the gene. Across the GTEx proteome ^18^, which is based on mass spectrometry rather than IHC, protein abundance falls from concordant through variable to suppressed genes. Secreted proteins, the group most likely to be lost during in situ staining, rank lowest by mass spectrometry rather than being recovered. Staining loss, therefore, does not account for the suppressed class. Randomness, the second possibility, would leave suppression unpredictable from sequence. Instead, a model trained on sequence-derived properties alone, with RNA withheld and no gene shared between training and test sets, predicts protein detection at an AUC of 0.755. Sequence has been shown to predict the protein to RNA ratio in bulk tissue ^20^. We show that sequence also predicts whether protein is detected at all, resolved to individual cell types. Suppression is therefore a genuine property of the gene, determined by its sequence rather than arising by chance.

The processes that uncouple protein from RNA have been characterized individually ^13^, and we resolve which of them produce the suppressed class. The deficit is primarily translational. Translational efficiency is reduced by roughly half in suppressed genes, while mRNA half-life is unchanged. This distinguishes them from the codon optimality pathway, in which poor translation and rapid transcript decay occur together ^8,9^. Suppression also arises after synthesis. A subunit that cannot assemble with its partners is degraded, and orphan subunits are among the largest sources of degraded protein in mammalian cells ^10,11^. In our atlas, the RNA rank of a subunit’s binding partners predicts its detection across most complexes, though the effect on any single subunit is modest. Assembly-dependent degradation therefore acts broadly across complexes, not only in the individual cases where it was first shown.

Human genetic evidence has become a primary basis for choosing drug targets, and targets with genetic support are roughly twice as likely to succeed in clinical development ^49^. This evidence supports two distinct steps, the identification of a candidate gene from disease and expression data, and the validation of that candidate by a compound shown to act on its protein ^50^. The suppressed class is enriched among targets nominated in DGIdb and Open Targets, which draw on clinical and literature evidence, and it is not enriched among targets in ChEMBL, which record measured compound activity. Identification uses RNA as a proxy for protein, and for the suppressed class RNA is not a reliable proxy. This is standard practice in neurodegeneration, where the brain is accessible only at autopsy. A protein found in accessible biofluid, such as CSF, is matched to a brain cell type using an RNA atlas and put forward as a brain-specific mechanism or drug target. From one RNA reading, researchers claim that the protein is present, that it comes from a specific cell type, and that it originates in the brain. RNA supports none of these claims because it can be abundant where protein is absent. Further, most protein in the lumbar CSF comes from blood rather than the brain ^46,48^. We built concordR and audited 3,304 CSF proteins nominated as brain cell type-enriched across this literature. Although most of these proteins were detectable in the brain, fewer than one in ten was specific to it and only 0.3% survived the full audit. The RNA to protein concordance in concordR is based on an aggregated estimate from across tissues. Some of the 318 brain-specific proteins, therefore, may be concordant too, however this requires further experimental follow-up. These findings highlight that RNA is a sufficient proxy for protein only in the concordant class. Claims about mechanism or drug targets in the suppressed and variable classes need protein-level confirmation before they are made.

## Limitations of the study

There are limitations to our study. First, RNA and protein come from separate donor cohorts, matched at the tissue and cell type level, because no genome-scale platform measures both in the same donor. However, each gene’s suppression rate aggregates its observations across all tissues and cell types, so donor mismatch has little effect on its class. We confirmed this by training a model on Tabula Sapiens and testing it on the separately collected Brain Cell Atlas, showing AUCs of 0.722 to 0.816. Second, IHC is antibody-based, and undetected protein could reflect antibody failure rather than true absence. We showed that using the antibody independent GTEx TMT-MS3 proteome reproduced the class ordering, minimizing the risk that our findings are an antibody artefact. Mass spectrometry still misses low abundance proteins, therefore future work should continue to develop single-cell proteomics and higher-affinity platforms ^51,52^. Third, the datasets we used to build the atlas are from healthy tissue because cell type-resolved datasets of disease tissue do not exist at genome scale. We note that suppression was predictable from sequence alone (AUC 0.755), and sequence does not change with disease, therefore gene classes should still hold across disease states. Further, we confirmed this by testing the proteins undetected in healthy brain against an Alzheimer’s disease cohort, recovering only three further proteins.

## Materials and Methods

### Data sources

Single-cell RNA expression data were obtained from Tabula Sapiens v2.0 ^22^, downloaded as organ level .h5ad files spanning 28 organs. Brain transcriptomics were sourced from the Human Cell Atlas Brain Cell Atlas v1.0 ^23,25^, as the brain is not represented in Tabula Sapiens. Protein level expression data were obtained from the Human Protein Atlas (HPA) normal tissue immunohistochemistry (IHC) dataset ^24^.

### Cross-dataset cell type harmonization

A consensus bridge map was manually curated to establish correspondence between HPA IHC cell type annotations and Tabula Sapiens cell ontology classes. Twenty-three organ level .h5ad files from Tabula Sapiens, spanning 868,136 cells were processed across 61 bridged cell types (1-6 cells per type; full organ x cell type breakdown in Supplementary Table 2). Log-normalized expression values from the log-normalized expression layer were used for all organs. Where a single HPA cell type encompasses functionally heterogeneous populations (e.g. pancreatic endocrine cells), it was mapped to multiple Tabula Sapiens cell types (11 such one-to-many mappings; maximum 4 Tabula Sapiens cell types per HPA cell type). These were subsequently aggregated at the HPA annotation level for RNA-protein comparisons.

For the brain, a separate bridge was constructed due to the distinct dataset source and the complexity of neuroanatomical nomenclature. Eighteen dissection-level .h5ad files from the Human Cell Atlas Brain Cell Atlas, spanning 813,870 cells, were mapped to five broad HPA-defined regions: caudate, cerebellum, cerebral cortex, hippocampus, and hypothalamus (1-6 dissections per region; full dissection-to-region mapping in Supplementary Table 3). Within each region, cells were assigned to one of two bins, neuronal or non-neuronal, and pseudobulk profiles were aggregated across sub-dissections by a cell count-weighted average.

### Pseudobulk RNA expression

For each Tabula Sapiens organ, per-cell-type pseudobulk expression profiles were computed by averaging single-cell expression values across all cells belonging to a given cell ontology class. Log-normalized expression values, stored in the log normalized layer, were used for all Tabula Sapiens organs. For the Brain Cell Atlas no pre-normalized layer was present, expression values were therefore library size normalized to 10,000 counts per cell and log(x+1)-transformed prior to pseudobulk averaging.

Tabula Sapiens and brain pseudobulk profiles were concatenated into a single expression matrix. A transcriptome-wide RNA expression percentile rank was then computed for each gene within each tissue x cell type stratum, using all detected genes as the denominator prior to any antibody coverage filtering. This ensures that rank reflects a gene’s position within the full expressed transcriptome rather than the subset with available protein data.

### Gene level feature extraction

Gene-level features were derived from three complementary sources. Transcript-level structural properties included transcript count, total transcript length, GC content and exon count were obtained from Ensembl BioMart (GRCh38) ^53^. Coding sequence features were computed from Ensembl cDNA FASTA sequences for 23,262 transcripts, including codon adaptation index (CAI), Kozak context score, 5′ and 3′ UTR lengths, 5′ and 3′ UTR CDS length, and 3′ UTR-to-CDS ratio. Protein-level biophysical features were derived from Ensembl canonical protein FASTA sequences (19,338 proteins); where multiple isoforms were present, the longest was retained. Features included protein length, molecular weight, global hydrophobicity and N-terminal hydrophobicity (mean over residues 1-25 and longest hydrophobic run within residues 1-30; Kyte-Doolittle scale ^54^), transmembrane helix count, length and residue fraction (hydrophobic segment proxy, threshold ≥ 18 residues), cysteine, charged, and proline residue fractions, N-glycosylation motif count (N-X-[S/T], X ≠ P), low-complexity fraction (sliding 20-residue window Shannon entropy, threshold < 2.2 bits), and intrinsic disorder fraction (Uversky charge-hydrophobicity heuristic, 25-residue window ^55^). Protein degradation propensity was characterized by four PEST motif features (maximum and summed PEST scores, number of qualifying regions, and fractional sequence coverage) computed using an implementation of the ePESTfind algorithm ^56–58^. After merging all sequence sources, 19,288 genes had a complete set of 30 sequence-derived features.

### External database features

Five external databases were integrated to capture post-transcriptional regulatory context and localization. Subcellular localization was obtained from UniProt (release 2026_01) where each protein was encoded across seven groups (secreted/vesicular, plasma membrane, nuclear, cytoplasmic, mitochondrial, ER/Golgi, cytoskeletal) plus a multi-localization flag. Protein complex membership was derived from CORUM 5.0 ^59^ core complexes (human entries only); features included binary membership, number of complexes, mean and maximum complex size, and a large-complex flag (≥10 subunits). miRNA regulation was obtained from TargetScan 8.0 ^60^ conserved site predictions. Features included number of conserved target sites, number of targeting miRNA families, and site density. Upstream open reading frame (uORF) features (count, maximum and mean uORF length, mean PhastCons conservation, and a presence flag) were derived from sORFs.org ^32^. RNA-binding protein occupancy (the total number of bound RBPs, binding density, and individual binding flags for HuR, hnRNP, and Pumilio) was quantified from ENCODE eCLIP data ^61^.

The combined gene-level feature set comprised of 59 features per gene. In the final quality filtered matrix, sequence-derived features achieved 98.7-98.9% coverage; external features ranged from 96.3% (subcellular localization binary flags, CORUM, TargetScan, sORF) to 98.9 % (ENCODE eCLIP).

### Master matrix construction and quality filtering

The master feature matrix was assembled by joining the combined RNA pseudobulk (4,723,322 rows; 61,856 unique ensembl entries, 28 tissues) with HPA IHC observations via the cell type bridge maps. HPA IHC expression levels labels “Descending”, “N/A”, “Not representative” as well as entries of “Uncertain” reliability were excluded prior to joining (1,199,675 raw rows reduced to 1,014,763 after reliability and level filtering). For brain tissues, 17,675 annotations (12.9%) describing neural processes rather than cell bodies (neuropil, synaptic glomeruli, dendrites, axonal projections) were excluded before IHC matching. After joining, 651,496 rows were obtained across both sources. Where multiple fine grained HPA cell types mapped to the same RNA cell type bin, rows were consolidated by retaining the highest protein score and strongest reliability tier; 87,757 duplicate groups (197,038 rows, 30.0% of pre-deduplication rows) were resolved, reducing the matrix to 542,215 rows.

The following sequential quality filters were then applied: (1) minimum pseudobulk cell count of 50 cells per stratum (removed 53,881 rows); (2) exclusion of tissues contributing fewer than 100 unique genes after filtering, which removed eye (26 genes), hypothalamus (39 genes), and ovary (20 genes). The final matrix comprised 488,190 observations across 11,154 unique genes, 24 tissues, and 53 cell types. One outcome variable was derived, which was a binary target (protein detected vs. not detected) with 261,291 positive (53.5%) and 226,899 negative (46.5%) observations.

### Gene classification and correlation decomposition

Gene-level classification was based on suppression rate, defined as the fraction of high RNA observations (RNA rank > 0.70) in which protein was not detected, computed for genes with at least five qualifying observations (n = 10,942). Four gene classes were defined by fixed thresholds: (1) suppressed (suppression rate > 0.80), (2) concordant (suppression rate < 0.20), (3) variable (0.20–0.80), and (4) low expression (fewer than 5 qualifying observations).

Spearman rank correlations between RNA expression percentile and protein detection score were computed at three levels, (1) globally across all observations, (2) within each gene discordance class (concordant, variable, suppressed), and (3) within each tissue and each tissue x cell type stratum. To assess whether the within-class correlation structure is conserved across tissues, correlations were computed separately for each gene class in each of the 24 tissues, yielding a gene class by tissue cross-decomposition matrix.

### GTEx mass spectrometry validation

To validate concordance classifications against an antibody independent platform, we used tissue specific quantitative mass spectrometry proteomics from the GTEx consortium ^18^. Median relative protein abundances (log_2_ transformed, normalized to a within-run reference) were obtained across 15 normal human tissue types, comprising tandem mass tag (TMT) 10-plex with MS3-level quantification of 12,627 genes from 14 donors. Gene identifiers were mapped to HGNC symbols ^62^ using BioMart mapping ^63,64^. For each gene, median protein abundance was calculated across all tissues in which it was quantified. Genes were then grouped by concordance class. The low expression class (n = 51 matched genes) was excluded owing to insufficient sample size. Differences in protein abundance between concordant and suppressed genes were tested using the Wilcoxon rank-sum test, and the relationship between protein confidence score and median protein abundance was assessed by Spearman rank correlation.

### Discordance modelling and gene classification

Binary protein detection was modelled using a histogram-based gradient-boosted tree classifier implemented in ‘scikit-learn’ ^65^, selected for its native handling of missing feature values at the scale of approximately 488,000 observations. Model inputs comprised RNA expression percentile rank and 58 gene-level features (29 sequence-derived, 29 from external databases). To prevent data leakage arising from the multi-observation-per-gene structure of the matrix, cross-validation was performed using five-fold GroupKFold with gene identity as the grouping variable, ensuring that all observations of a given gene were confined to either the training or test set within each fold. Out-of-fold (OOF) predictions were collected to produce an unbiased probability estimate for every observation. Hyperparameters were tuned using nested cross-validation. Within each outer training fold, a three-fold inner gene grouped cross-validation evaluated 30 randomly sampled configurations, selecting the combination that maximized ROC-AUC. The search space covered learning rate, number of boosting iterations, maximum tree depth, minimum leaf size, and L2 regularization strength. Optimal hyperparameters from each outer fold were aggregated into a single consensus configuration, which was used for all reported analyses. This approach avoids fold-to-fold variability in model specification while preserving the leakage prevention guarantees of the nested design. Model calibration was assessed via expected calibration error (ECE, 10 equal width bins) on OOF predictions.

### Cross-source validation

To evaluate cross-platform generalization, the detection model was retrained on all Tabula Sapiens observations and evaluated on Brain Cell Atlas observations without any fine-tuning or domain adaptation. This constitutes a strict cross-source test. Training and test data were derived from independent datasets, collected on different platforms (10x Chromium for Tabula Sapiens; Smart-seq2 for Brain Cell Atlas), and processed through separate pseudobulk pipelines. Gene-level protein confidence scores (suppression rates) computed from the Tabula training set were correlated with per-gene brain detection rates to assess cross-source transfer of gene-level discordance profiles. A within-Tabula 5-fold gene-grouped CV baseline was computed on the same model for comparison.

### Molecular determinants of translational fate

To characterize the molecular determinants of translational fate, univariate comparisons between consistently concordant and consistently suppressed genes were conducted across all 58 features using Cohen’s d as the effect size measure, with false discovery rate (FDR)-corrected Wilcoxon rank-sum tests for statistical significance.

To test whether the separation between concordant and suppressed genes is additive or requires interactions among features, we compared two model classes on the concordant-versus-suppressed task. They were both restricted to the 15 features with the largest absolute Cohen’s d: a logistic regression, which is purely additive, and a gradient-boosted tree classifier, which can capture feature interactions. Both were implemented in “scikit-learn” and evaluated under an identical protocol, the same 5-fold gene-grouped cross-validation (GroupKFold on Ensembl gene identity) and, within each training fold, the same reselection of the top 15 features by absolute Cohen’s d. For the logistic model, features were additionally standardized to zero mean and unit variance, and the model was L2-regularised (C = 1.0, maximum iterations = 1000). AUCs are reported as the mean ± standard deviation across folds.

### CORUM stoichiometry test

To test whether protein complex membership contributes to tissue-specific protein detection patterns through stoichiometric quality control, we assessed whether a subunit’s protein detection across tissues correlates with the RNA expression of the other subunits in the same complex. Human protein complexes with > 3 subunits were obtained from CORUM v5.0 ^59^ (2,599 complexes, 3,974 unique genes). For each focal gene in a complex, partner RNA co-expression was computed as the mean RNA expression percentile of all other subunits in the same complex, calculated separately for each of the 24 tissues. The Spearman correlation between partner co-expression and focal gene protein detection was then computed across tissues for each gene-complex pair, yielding a stoichiometry coefficient ρ_stoich_. After filtering for atlas coverage and sufficient between-tissue variance, 1,502 complexes (6,716 gene-complex pairs covering 2,115 unique focal genes) were retained for analysis. Statistical significance was assessed three ways: (1) a one sample t-test of ρ_stoich_ against zero across all gene-complex pairs, (2) the fraction of pairs with positive ρ_stoich_, and (3) a permutation control (500 iterations) in which partner gene identities were shuffled to generate a null distribution for the expected mean ρ_stoich_.

### Translational efficiency

Translational efficiency (TE) estimates were obtained from RPFdb v3.0 ^31^, a compendium of ribosome profiling studies. Per-gene TE values, defined as ribosome-protected fragment density normalized to mRNA abundance, were extracted from the H. sapiens coding sequence (CDS)-level archive. Ensembl gene identifiers were stripped of version suffixes, and a single median TE was computed per gene across all available samples. TE distributions were compared across the three discordance classes (consistently concordant, variable, consistently suppressed) using the Mann-Whitney U test, with Cohen’s d as an effect size measure.

### Protein and mRNA half-life integration

To identify the molecular mechanism(s) leading to protein suppression, protein and mRNA turnover were compared across the three gene classes (concordant, variable, suppressed). Only genes assigned to these classes in the discordance atlas were retained.

#### Protein half-life

Proteome-wide protein half-lives were obtained from Mathieson et al. ^12^, restricted to the human primary cell types assayed (B cells, NK cells, hepatocytes, monocytes). Mouse measurements were excluded. For each gene, half-life values were collected across all human cell-type columns for which both the value and its paired per-measurement quality annotation were present and numeric. A gene-level estimate was taken as the median across all valid human measurements, and, separately, as the median across measurements flagged “good” quality, the latter used for a sensitivity analysis. Half-lives were log_2_-transformed for all statistical tests and back-transformed to hours for reporting. Gene-level values were joined to the atlas classes by gene symbol.

#### mRNA half-life

mRNA half-lives were obtained from RNADecayCafe ^30^, a uniformly reprocessed dataset of SLAM-seq/TimeLapse-seq decay measurements across human cell lines, re-quantified with EZbakR. The donor normalized half-life estimate was used, as it is better suited to cross-dataset comparison. Measurements were filtered to expressed transcripts (RPKM > 0 in either the total or 3′-end assay) and to positive, non-missing half-lives. Per-gene summaries were computed as the median (with mean and standard deviation) across all cell lines, and additionally as the median restricted to four hematopoietic lines (K562, MOLM-13, MV4-11, Nalm-6) for a cell type-matched sensitivity analysis.

For both protein and mRNA half-life, differences in log_2_ half-life among the three classes were assessed by the Kruskal-Wallis test, followed by pairwise two-sided Mann-Whitney U tests (concordant vs. suppressed, concordant vs. variable, variable vs. suppressed) with Cohen’s d as the effect-size measure (positive values denoting longer half-life in concordant genes). The monotonic association between per-gene suppression rate and log□ half-life was quantified with Spearman’s ρ.

### Drug target enrichment analysis

Drug target annotations were obtained from two sources. ChEMBL v34 ^33^ entries were filtered to single-protein targets (9,248 entries; 4,246 unique human genes), excluding multi-protein complexes and protein families. Interactions from the Drug Gene Interaction Database (DGIdb) v5.0 ^34^ were filtered to medium-to-high confidence (score > 0.5), retaining 26,035 of 98,239 interactions across 4,081 unique genes, of which 3,244 were annotated as approved drug targets. Target tractability annotations were obtained from the Open Targets Platform ^35^ release 24.09, downloading the dataset as partitioned Parquet files from the EMBL-EBI FTP server (https://ftp.ebi.ac.uk/pub/databases/opentargets/platform/24.09/). The 200 files were consolidated into a single table retaining gene identifier, tractability, and target class. From the tractability annotations we extracted clinical precedence flags per modality (small molecule, antibody, PROTAC, other), defining a gene as clinically tractable if any modality carried an “Approved Drug”, “Advanced Clinical”, or “Phase 1 Clinical” assessment. Enrichment of each annotation set across concordance classes was assessed using Fisher’s exact test against a background of all atlas genes. Odds ratios are reported with Benjamini-Hochberg-corrected *p*-values. ChEMBL and DGIdb were analysed separately, as they are biochemical assay data and pharmacogenomic annotations, respectively.

### Biomarker and drug target assessment

To evaluate whether atlas-derived concordance classifications can inform the interpretation of biofluid proteomics findings, we applied the discordance atlas to differentially abundant proteins (DAPs) reported as being brain cell-derived based on RNA evidence across several published cerebrospinal fluid (CSF) studies spanning neurodegenerative diseases including Alzheimer’s disease, frontotemporal dementia, Lewy Body dementia, Parkinson’s disease, and amyotrophic lateral sclerosis ^36–44^. These studies employed various proteomic platforms (SomaScan 7k assay, Olink proximity extension assay (PEA), and TMT mass spectrometry), providing a broad test of platform-independent applicability. DAPs were collated into a CSF (3,304 proteins) and plasma (16 proteins) panel. Each protein was annotated with its concordance class, mechanistic tier, subcellular localization, tissue specificity, and a composite reliability assessment reflecting whether its protein presence and level are consistently predicted by RNA abundance across tissues. Panel coverage and annotation distributions were tabulated for both CSF and plasma.

To assess whether concordance flags assigned to audited proteins reflect genuine biological absence rather than artefacts of HPA IHC, we validated against human brain tissue using TMT mass spectrometry data from the Accelerating Medicines Partnership-Alzheimer’s Disease (AMP-AD) Diverse cohorts project (1,275 samples; 8,435 quantified proteins) restricting to Alzheimer’s disease (n = 876) and cognitively normal control (n = 399) samples ^45^. Only proteins with < 30% missing values across the sample set were retained. The remaining missing values were imputed using per-protein median values. Residual batch effects were corrected by fitting a linear regression model, consistent with the approach used in the original study ^45^.

### ConcordR

The atlas is implemented in concordR (https://github.com/Art83/concordR), an R package that operationalizes it for translational research evaluating proteins from proteomics studies. It bundles the pre-computed concordance data (11,154 genes, 24 tissues, 53 cell types) so that audits run without external dependencies. Given a gene list and a sample type (CSF, plasma, or urine) with an optional tissue context, its triage() function applies four sequential criteria: (1) detection of protein in brain, (2) a physical route from brain to biofluid, (3) specificity of the protein to brain, and (4) concordance between RNA and protein. To avoid antibody-specific blind spots, protein-level detection does not rest on IHC alone. A gene is counted as detected in a tissue if supported by any of three orthogonal sources, HPA IHC, GTEx mass spectrometry, or PaxDb mass spectrometry, so a tissue-of-origin mismatch is called only when a protein is absent across both antibody- and peptide-based platforms. We assess the physical route from annotated subcellular localization. An intracellular protein (e.g. in mitochondrion, Golgi Apparatus, endoplasmic reticulum, etc.) has no path to a biofluid through the secretory pathway or proteolytic cleavage and is deemed implausible. We do not classify the route as implausible in instances where the protein also carries a neural projection annotation (e.g. axon, dendrite, synapse, axon terminal, neuron projection). Neuronal projections release their contents into the extracellular space through synaptic turnover and axonal degeneration, and the released proteins enter CSF. These proteins are flagged in concordR for the user. Each gene is returned with a verdict and the per-criterion evidence behind it, leaving the user to define which verdicts constitute failures for their specific claim. Companion functions (query_atlas, query_tissue, plot_gene) expose the underlying per-gene and tissue-resolved data directly.

## Data Availability

The current work used publicly available data. All data generated in this work is available at Zenodo under DOI https://doi.org/10.5281/zenodo.21273384.

## Code availability

All code used in this manuscript is publicly available at https://github.com/Art83/hpa_to_tabula. The concordR R package is available at https://github.com/Art83/concordR.

## Author Contributions

Data Curation, Formal Analysis, Investigation, Methodology, Software, Resources, Visualization: A.S.

Conceptualization, Funding acquisition, Writing – Original Draft, Writing – Review & Editing: C.A.F. and A.S.

## Supplementary Figure

**Supplementary Figure 1.**
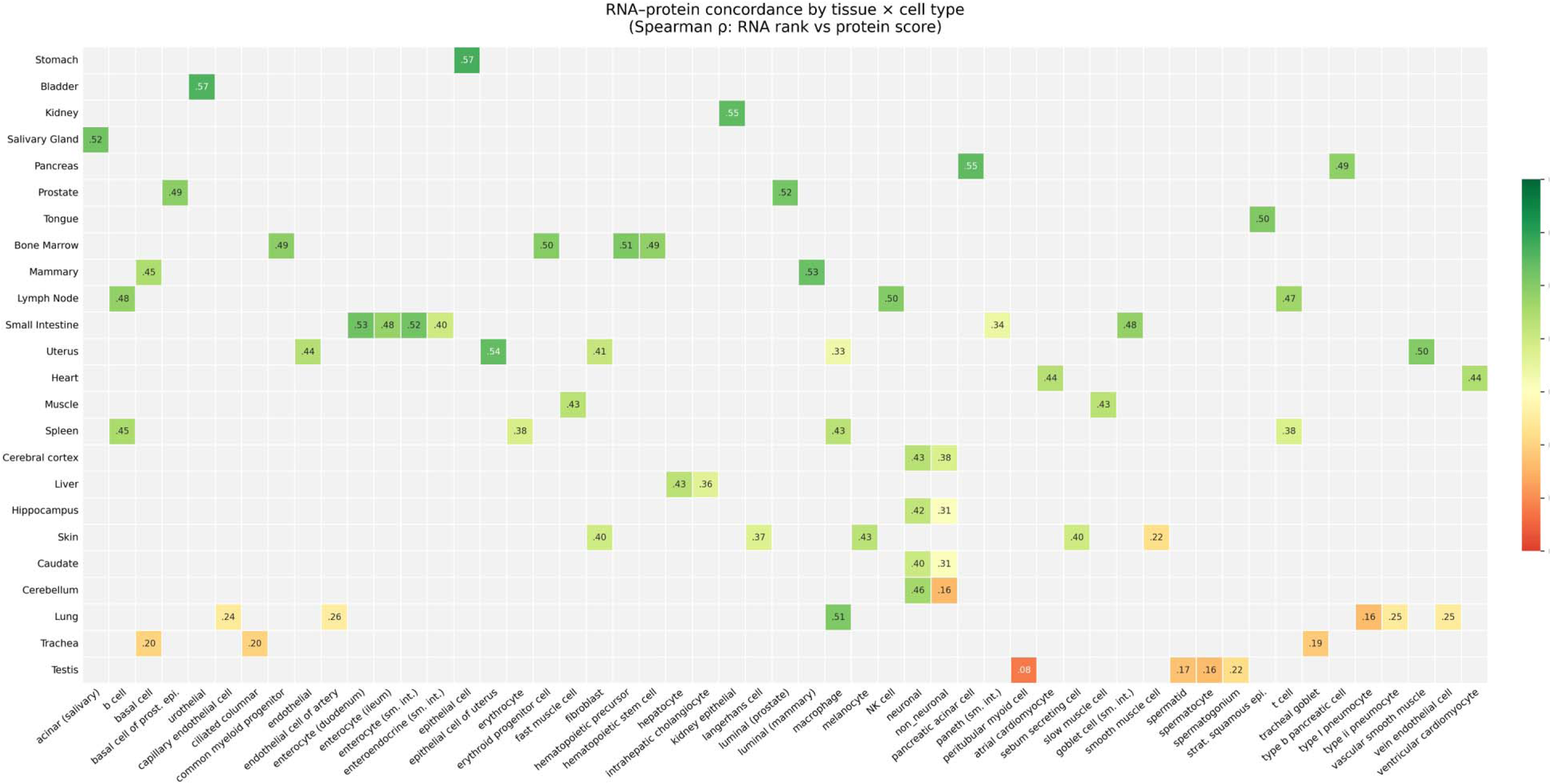
RNA to protein concordance varies across cell types. Spearman correlation between RNA rank and protein detection within each cell type. Cell types are ordered from highest to lowest correlation and colored by tissue of origin. Values ranged from ρ = 0.572 in stomach epithelial cells to ρ = 0.077 in testis peritubular myoid cells.

## References

1. Schwanhausser, B., Busse, D., Li, N., Dittmar, G., Schuchhardt, J., Wolf, J., Chen, W., and Selbach, M. (2011). Global quantification of mammalian gene expression control. Nature 473, 337–342.

2. Vogel, C., and Marcotte, E.M. (2012). Insights into the regulation of protein abundance from proteomic and transcriptomic analyses. Nature Reviews Genetics 13, 227–232.

3. Liu, Y., Beyer, A., and Aebersold, R. (2016). On the dependency of cellular protein levels on mRNA abundance. Cell 165, 535–550.

4. de Sousa Abreu, R., Penalva, L.O., Marcotte, E.M., and Vogel, C. (2009). Global signatures of protein and mRNA expression levels. Molecular bioSystems 5, 1512–1526.

5. Baek, D., Villen, J., Shin, C., Camargo, F.D., Gygi, S.P., and Bartel, D.P. (2008). The impact of microRNAs on protein output. Nature 455, 64–71.

6. Guo, H., Ingolia, N.T., Weissman, J.S., and Bartel, D.P. (2010). Mammalian microRNAs predominantly act to decrease target mRNA levels. Nature 466, 835–840.

7. Selbach, M., Schwanhausser, B., Thierfelder, N., Fang, Z., Khanin, R., and Rajewsky, N. (2008). Widespread changes in protein synthesis induced by microRNAs. Nature 455, 58–63.

8. Hanson, G., and Coller, J. (2018). Codon optimality, bias and usage in translation and mRNA decay. Nature Reviews Molecular Cell Biology 19, 20–30.

9. Presnyak, V., Alhusaini, N., Chen, Y.-H., Martin, S., Morris, N., Kline, N., Olson, S., Weinberg, D., Baker, K.E., Graveley, B.R., and Coller, J. (2015). Codon optimality is a major determinant of mRNA stability. Cell 160, 1111–1124.

10. Juszkiewicz, S., and Hegde, R.S. (2018). Quality control of orphaned proteins. Molecular Cell 71, 443–457.

11. McShane, E., Sin, C., Zauber, H., Wells, J.N., Donnelly, N., Wang, X., Hou, J., Chen, W., Storchova, Z., Marsh, J.A., et al. (2016). Kinetic analysis of protein stability reveals age-dependent degradation. Cell 167, 803–815.

12. Mathieson, T., Franken, H., Kosinski, J., Kurzawa, N., Zinn, N., Sweetman, G., Poeckel, D., Ratnu, V.S., Schramm, M., Becher, I., et al. (2018). Systematic analysis of protein turnover in primary cells. Nature Communications 9, 689.

13. Buccitelli, C., and Selbach, M. (2020). mRNAs, proteins and the emerging principles of gene expression control. Nature Reviews Genetics 21, 630–644.

14. Gygi, S.P., Rochon, Y., Franza, B.R., and Aebersold, R. (1999). Correlation between protein and mRNA abundance in yeast. Molecular and Cellular Biology 19, 1720–1730.

15. Maier, T., Guell, M., and Serrano, L. (2009). Correlation of mRNA and protein in complex biological samples. FEBS Letters 583, 3966–3973.

16. Edfors, F., Danielsson, F., Hallstrom, B.M., Kall, L., Lundberg, E., Ponten, F., Forsstrom, B., and Uhlen, M. (2016). Gene-specific correlation of RNA and protein levels in human cells and tissues. Molecular Systems Biology 12, 883.

17. Franks, A., Airoldi, E., and Slavov, N. (2017). Post-transcriptional regulation across human tissues. PLOS Computational Biology 13, e1005535.

18. Jiang, L., Wang, M., Lin, S., Jian, R., Li, X., Chan, J., Dong, G., Fang, H., Robinson, A.E., GTEx Consortium, and Snyder, M.P. (2020). A quantitative proteome map of the human body. Cell 183, 269–283.

19. Wang, D., Eraslan, B., Wieland, T., Hallstrom, B.M., Hopf, T., Zolg, D.P., Zecha, J., Asplund, A., Li, L.-H., Meng, C., et al. (2019). A deep proteome and transcriptome abundance atlas of 29 healthy human tissues. Molecular Systems Biology 15, e8503.

20. Eraslan, B., Wang, D., Gusic, M., Prokisch, H., Hallstrom, B.M., Uhlen, M., Asplund, A., Ponten, F., Wieland, T., Hopf, T., et al. (2019). Quantification and discovery of sequence determinants of protein-per-mRNA amount in 29 human tissues. Molecular Systems Biology 15, e8513.

21. Karlsson, M., Zhang, C., Mear, L., Zhong, W., Digre, A., Katona, B., Sjostedt, E., Butler, L., Odeberg, J., Dusart, P., et al. (2021). A single-cell transcriptomics map of human tissues. Science Advances 7, eabh2169.

22. Tabula Sapiens Consortium (2022). The Tabula Sapiens: A multiple-organ, single-cell transcriptomic atlas of humans. Science 376, eabl4896.

23. Siletti, K., Hodge, R., Albiach, A.M., Lee, K.W., Ding, S.-L., Hu, L., Lonnerberg, P., Bakken, T., Casper, T., Clark, M., et al. (2023). Transcriptomic diversity of cell types across the adult human brain. Science 382, eadd7046.

24. Uhlen, M., Fagerberg, L., Hallstrom, B.M., Lindskog, C., Oksvold, P., Mardinoglu, A., Sivertsson, A., Kampf, C., Sjostedt, E., Asplund, A., et al. (2015). Tissue-based map of the human proteome. Science 347, 1260419.

25. Regev, A., Teichmann, S.A., Lander, E.S., Amit, I., Benoist, C., Birney, E., Bodenmiller, B., Campbell, P., Carninci, P., Clatworthy, M., et al. (2017). The Human Cell Atlas. Elife 6.

26. Soumillon, M., Necsulea, A., Weier, M., Brawand, D., Zhang, X., Gu, H., Barthes, P., Kokkinaki, M., Nef, S., Gnirke, A., et al. (2013). Cellular source and mechanisms of high transcriptome complexity in the mammalian testis. Cell Reports 3, 2179–2190.

27. Kleene, K.C. (2003). Patterns, mechanisms, and functions of translation regulation in mammalian spermatogenic cells. Cytogenetic and Genome Research 103, 214–224.

28. Gou, L.-T., Dai, P., Yang, J.-H., Xue, Y., Hu, Y.-P., Zhou, Y., Kang, J.-Y., Wang, X., Li, H., Hua, M.-M., et al. (2014). Pachytene piRNAs instruct massive mRNA elimination during late spermiogenesis. Cell Research 24, 680–700.

29. Hilz, S., Modzelewski, A.J., Cohen, P.E., and Grimson, A. (2016). The roles of microRNAs and siRNAs in mammalian spermatogenesis. Development 143, 3061–3073.

30. Vock, I.W., Tang, D., Giraldez, A.J., and Simon, M.D. (2025). RNADecayCafe, a uniformly processed atlas of RNA half-life estimates across multiple human cell lines. bioRxiv.

31. Wang, Y., Tang, Y., Xie, Z., and Wang, H. (2025). RPFdb v3.0: An enhanced repository for ribosome profiling data and related content. Nucleic Acids Research 53, D293–D298.

32. Olexiouk, V., Crappe, J., Verbruggen, S., Verhegen, K., Martens, L., and Menschaert, G. (2016). sORFs.org: A repository of small ORFs identified by ribosome profiling. Nucleic Acids Research 44, D324–D329.

33. Mendez, D., Gaulton, A., Bento, A.P., Chambers, J., De Veij, M., Felix, E., Magarinos, M.P., Mosquera, J.F., Mutowo, P., Nowotka, M., et al. (2019). ChEMBL: Towards direct deposition of bioassay data. Nucleic Acids Research 47, D930–D940.

34. Cannon, M., Stevenson, J., Stahl, K., Basu, R., Coffman, A., Kiwala, S., McMichael, J.F., Kuzma, K., Morrissey, D., Cotto, K., et al. (2024). DGIdb 5.0: Rebuilding the drug-gene interaction database for precision medicine and drug discovery platforms. Nucleic Acids Research 52, D1227–D1235.

35. Koscielny, G., An, P., Carvalho-Silva, D., Cham, J.A., Fumis, L., Gasparyan, R., Hasan, S., Karamanis, N., Maguire, M., Papa, E., et al. (2016). Open Targets: A platform for therapeutic target identification and validation. Nucleic Acids Research 45, D985–D994.

36. Ali, M., Timsina, J., Western, D., Liu, M., Beric, A., Budde, J., Do, A., Heo, G., Wang, L., Gentsch, J., et al. (2025). Multi-cohort cerebrospinal fluid proteomics identifies robust molecular signatures across the Alzheimer disease continuum. Neuron 113, 1363–1379.

37. Ali, M., Timsina, J., Xu, Y., Chen, Y., Gong, K., Western, D., Heo, G., Liu, M., Budde, J., Pottier, C., et al. (2026). Large-scale CSF and plasma proteomics reveal immune, synaptic, and extracellular matrix disruptions across neurodegenerative diseases. Neuron.

38. Binette, A.P., Gaiteri, C., Wennstrom, M., Kumar, A., Hristovska, I., Spotorno, N., Salvado, G., Strandberg, O., Mathys, H., Tsai, L.-H., et al. (2024). Proteomic changes in Alzheimer’s disease associated with progressive Aβ plaque and tau tangle pathologies Nature Neuroscience 27, 1880–1891.

39. Tijms, B.E., Vromen, E.M., Mjaavatten, O., Holstege, H., Reus, L.M., van der Lee, S., Wesenhagen, K.E.J., Lorenzini, L., Vermunt, L., Venkatraghavan, V., et al. (2024). Cerebrospinal fluid proteomics in patients with Alzheimer’s disease reveals five molecular subtypes with distinct genetic risk profiles. Nature Aging 4, 33–47.

40. Blujdea, E.-R., van Bokhoven, P., Martino-Adami, P.V., Marshe, V.S., Vromen, E.M., Hok-A-Hin, Y.S., Boiten, W.A., Irwin, D.J., Chen-Plotkin, A.S., Lemstra, A.W., et al. (2026). Microglia protein profiles in CSF across Alzheimer’s disease clinical stages. Nature Aging 6, 520–533.

41. Johnson, E.C.B., Dammer, E.B., Duong, D.M., Ping, L., Zhou, M., Yin, L., Higginbotham, L.A., Guajardo, A., White, B., Troncoso, J.C., et al. (2020). Large-scale proteomic analysis of Alzheimer’s disease brain and cerebrospinal fluid reveals early changes in energy metabolism associated with microglia and astrocyte activation. Nature Medicine 26, 769–780.

42. Trautwig, A.N., Fox, E.J., Dammer, E.B., Shantaraman, A., Ping, L., Duong, D.M., Watson, C.M., Wu, F., Asress, S., Guo, Q., et al. (2025). Network analysis of the cerebrospinal fluid proteome reveals shared and unique differences between sporadic and familial forms of amyotrophic lateral sclerosis. Molecular Neurodegeneration 20, 58.

43. Shen, Y., Timsina, J., Heo, G., Beric, A., Ali, M., Wang, C., Yang, C., Wang, Y., Western, D., Liu, M., et al. (2024). CSF proteomics identifies early changes in autosomal dominant Alzheimer’s disease. Cell 187, 6309–6326.

44. Modeste, E.S., Ping, L., Watson, C.M., Duong, D.M., Dammer, E.B., Johnson, E.C.B., Roberts, B.R., Lah, J.J., Levey, A.I., and Seyfried, N.T. (2023). Quantitative proteomics of cerebrospinal fluid from African Americans and Caucasians reveals shared and divergent changes in Alzheimer’s disease. Molecular Neurodegeneration 18, 48.

45. Seifar, F., Fox, E.J., Shantaraman, A., Liu, Y., Dammer, E.B., Modeste, E.S., Duong, D.M., Yin, L., Trautwig, A.N., Guo, Q., et al. (2024). Large-scale deep proteomic analysis in Alzheimer’s disease brain regions across race and ethnicity. Alzheimer’s & Dementia 20, 8878–8897.

46. Sakka, L., Coll, G., and Chazal, J. (2011). Anatomy and physiology of cerebrospinal fluid. European Annals of Otorhinolaryngology, Head and Neck Diseases 128, 309–316.

47. Regeniter, A., Kuhle, J., Mehling, M., Moller, H., Wurster, U., Freidank, H., and Siede, W.H. (2009). A modern approach to CSF analysis: Pathophysiology, clinical application, proof of concept and laboratory reporting. Clinical Neurology and Neurosurgery 111, 313–318.

48. Reiber, H. (2001). Dynamics of brain-derived proteins in cerebrospinal fluid. Clinica Chimica Acta 310, 173–186.

49. Nelson, M.R., Tipney, H., Painter, J.L., Shen, J., Nicoletti, P., Shen, Y., Floratos, A., Sham, P.C., Li, M.J., Wang, J., et al. (2015). The support of human genetic evidence for approved drug indications. Nature Genetics 47, 856–860.

50. Finan, C., Gaulton, A., Kruger, F.A., Lumbers, R.T., Shah, T., Engmann, J., Galver, L., Kelley, R., Karlsson, A., Santos, R., et al. (2017). The druggable genome and support for target identification and validation in drug development. Science Translational Medicine 9.

51. Bennett, H.M., Stephenson, W., Rose, C.M., and Darmanis, S. (2023). Single-cell proteomics enabled by next-generation sequencing or mass spectrometry. Nature Methods 20, 363–374.

52. Kelly, R.T. (2020). Single-cell proteomics: Progress and prospects. Molecular & Cellular Proteomics 19, 1739–1748.

53. Smedley, D., Haider, S., Ballester, B., Holland, R., London, D., Thorisson, G., and Kasprzyk, A. (2009). BioMart -Biological queries made easy. BMC Genomics 10.

54. Kyte, J., and Doolittle, R.F. (1982). A simple method for displaying the hydropathic character of a protein. Journal of Molecular Biology 157, 105–132.

55. Uversky, V.N., Gillespie, J.R., and Fink, A.L. (2000). Why are “natively unfolded” proteins unstructured under physiologic conditions? Proteins 41, 415–427.

56. Rogers, S., Wells, R., and Rechsteiner, M. (1986). Amino acid sequences common to rapidly degraded proteins: The PEST hypothesis. Science 234, 364–368.

57. Rechsteiner, M., and Rogers, S. (1996). PEST sequences and regulation by proteolysis. Trends in Biochemical Sciences 21, 267–271.

58. Rice, P., Longden, I., and Bleasby, A. (2000). EMBOSS: The European Molecular Biology Open Software Suite. Trends in Genetics 16, 276–277.

59. Steinkamp, R., Tsitsiridis, G., Brauner, B., Montrone, C., Fobo, G., Frishman, G., Avram, S., Oprea, T.I., and Ruepp, A. (2024). CORUM in 2024: Protein complexes as drug targets. Nucleic Acids Research 53, D651–D657.

60. McGeary, S.E., Lin, K.S., Shi, C.Y., Pham, T.M., Bisaria, N., Kelley, G.M., and Bartel, D.P. (2019). The biochemical basis of microRNA targeting efficacy. Science 366.

61. Van Nostrand, E.L., Freese, P., Pratt, G.A., Wang, X., Wei, X., Xiao, R., Blue, S.M., Chen, J.-Y., Cody, N.A.L., Dominguez, D., et al. (2020). A large-scale binding and functional map of human RNA-binding proteins. Nature 583, 711–719.

62. Seal, R.L., Braschi, B., Gray, K., McClay, J., Tweedie, S., and Bruford, E.A. (2026). Genenames.org: The HGNC and PGNC resources in 2026. Nucleic Acids Research 54, D1098–D1107.

63. Durinck, S., Moreau, Y., Kasprzyk, A., Davis, S., De Moor, B., Brazma, A., and Huber, W. (2005). BioMart and Bioconductor: A powerful link between biological databases and microarray data analysis. Bioinformatics 21, 3439–3440.

64. Durinck, S., Spellman, P.T., Birney, E., and Huber, W. (2009). Mapping identifiers for the integration of genomic datasets with the R/Bioconductor package biomaRt. Nature Protocols 4, 1184–1191.

65. Pedregosa, F., Varoquaux, G., Gramfort, A., Michel, V., Thirion, B., Grisel, O., Blondel, M., Prettenhofer, P., Weiss, R., Dubourg, V., et al. (2011). Scikit-learn: Machine learning in Python. Journal of Machine Learning Research 12, 2825–2830.

